# Integrating phylogenetics with intron positions illuminates the origin of the complex spliceosome

**DOI:** 10.1101/2022.08.31.505394

**Authors:** Julian Vosseberg, Daan Stolker, Samuel H. A. von der Dunk, Berend Snel

## Abstract

Eukaryotic genes are characterised by the presence of introns that are removed from the pre-mRNA by the spliceosome. This ribonucleoprotein complex is comprised of multiple RNA molecules and over a hundred proteins, which makes it one of the most complex molecular machines that originated during the prokaryote-to-eukaryote transition. Previous work has established that these introns and the spliceosomal core originated from self-splicing introns in prokaryotes. Yet it remains largely elusive how the spliceosomal core expanded by recruiting many additional proteins. In this study we use phylogenetic analyses to infer the evolutionary history of the 145 proteins that we could trace back to the spliceosome in the last eukaryotic common ancestor (LECA). We found that an overabundance of proteins derived from ribosome-related processes were added to the prokaryote-derived core. Extensive duplications of these proteins substantially increased the complexity of the emerging spliceosome. By comparing the intron positions between spliceosomal paralogs, we infer that most spliceosomal complexity postdates the spread of introns through the proto-eukaryotic genome. The reconstruction of early spliceosomal evolution provides insight into the driving forces behind the emergence of complexes with many proteins during eukaryogenesis.

## Introduction

The spliceosome is a dynamic ribonucleoprotein (RNP) complex that assembles on the pre-mRNA to remove introns, intervening sequences between the exons. The exons are spliced together to form mature mRNA. Like the complex, the exon-intron structure of protein-coding genes is characteristic of eukaryotes. Transcription and splicing occur in the nucleus, which physically separates these processes from protein translation. Failure of correct splicing generally results in non-functional proteins.

The composition of the spliceosome changes during the splicing cycle (Wilkinson et al. 2020). It consists of the five small nuclear RNAs (snRNAs) U1, U2, U4, U5 and U6, which are bound by multiple proteins to form small nuclear RNPs (snRNPs), and several additional subcomplexes and factors. In the splicing reaction, the 5’ splice site first reacts with the adenosine branch point, forming a lariat structure. Subsequently, the exons are ligated and the lariat intron is released. The components of the spliceosome orchestrate different activities in a precisely ordered manner: they recognise the splice sites and the branch point sequences, prevent a premature reaction, perform the splicing reaction and assemble, remodel or disassemble the complex. The spliceosome is one of the most complex molecular machineries in eukaryotic cells and a complex spliceosome was present in the last eukaryotic common ancestor (LECA) (Collins and Penny 2005).

Eukaryotes have two types of introns that are recognised by different spliceosome complexes. The vast majority of introns are of U2-type and are recognised by the major spliceosome; U12-type introns comprise a small minority (Moyer et al. 2020). The minor spliceosome specifically recognises U12-type introns and most proteins of the major spliceosome are also part of the minor spliceosome (Turunen et al. 2013; Bai et al. 2021). All snRNAs but U5 have a minor-spliceosome equivalent (U11, U12, U4atac and U6atac) and a few minor-spliceosome specific proteins have been identified, especially in the U11/U12 di-snRNP (Turunen et al. 2013). The minor spliceosome and U12-type introns were also present in LECA (Russell et al. 2006).

In sharp contrast to a probably intron-rich LECA (Csuros et al. 2011; Vosseberg et al. 2022) with a complex spliceosome, prokaryotes lack intragenic introns and a spliceosome, meaning that they must have emerged at some time during eukaryogenesis. Spliceosomal introns and the key spliceosomal protein PRPF8 are thought to derive from self-splicing group II introns in prokaryotic genomes. This is based on similarities in the splicing reaction, function and structure of the RNAs involved, as well as the homology inferred between the spliceosomal protein PRPF8 and the single protein encoded by group II introns, the intron-encoded protein (IEP) (Zimmerly and Semper 2015). Recent work has suggested that the emergence of intragenic introns might have been an early event during eukaryogenesis (Vosseberg et al. 2022). The evolutionary histories of a few gene families in the spliceosome have been described (Anantharaman et al. 2002; Veretnik et al. 2009; Califice et al. 2012) and they suggest gene duplications played a pivotal role in the emergence of the complex spliceosome. Yet, a detailed picture of the origins of the full spliceosome, one of the most complex machineries to emerge during eukaryogenesis, is lacking.

This paper details in-depth phylogenetic analyses to reconstruct the spliceosome in LECA and the evolutionary histories of these LECA proteins in the prokaryote-to-eukaryote transition. Subsequent integration of the phylogenetic trees with the positions of introns allows to investigate the relation between the origin of the spliceosome and the emergence of intragenic introns. Our findings underline the role of gene duplications in establishing the complex LECA spliceosome and we detected a strong evolutionary link with the ribosome. The intron analyses suggest that the emergence of a complex spliceosome occurred late relative to the spread of introns.

## Results

### Complex composition of the LECA spliceosome

To infer the evolutionary origin of the LECA spliceosome, it is first necessary to establish which proteins were likely present in the spliceosome in LECA. The most recent systematic inventory of the composition of LECA’s spliceosome stems from 2005 (Collins and Penny 2005) and since then multiple additional proteins, such as the minor-spliceosome specific proteins, have been traced back to the eukaryotic ancestor. In conjunction with the enormous increase in genomic data, this provides ample reasons to update the reconstruction of the composition of the LECA spliceosome. We carried out this reconstruction by performing homology searches with spliceosomal proteins of human (Supplementary Table 1) and baker’s yeast (Supplementary Table 2), two species whose spliceosomes are well-studied. We used a strict definition of the spliceosome, which excludes proteins that function in related processes such as the coupling of splicing with transcription and the regulation of splicing. 145 spliceosomal orthogroups (OGs) could be traced to LECA (Figure 1, Supplementary Table 3). This number is nearly twice as large as the previously estimated 78 spliceosomal proteins in LECA (Collins and Penny 2005), a consequence of the expanded genomic sampling of eukaryotic biodiversity and increased knowledge on eukaryotic spliceosomes. The inferred number of spliceosomal LECA OGs is slightly lower than the number of spliceosomal proteins in human (164, only one LECA OG missing) and substantially larger than the number of proteins in the yeast spliceosome (99, 86 LECA OGs present). In addition to these proteins, five major spliceosomal snRNAs and the four minor-spliceosome specific snRNAs were also present in LECA.

**Figure 1.**
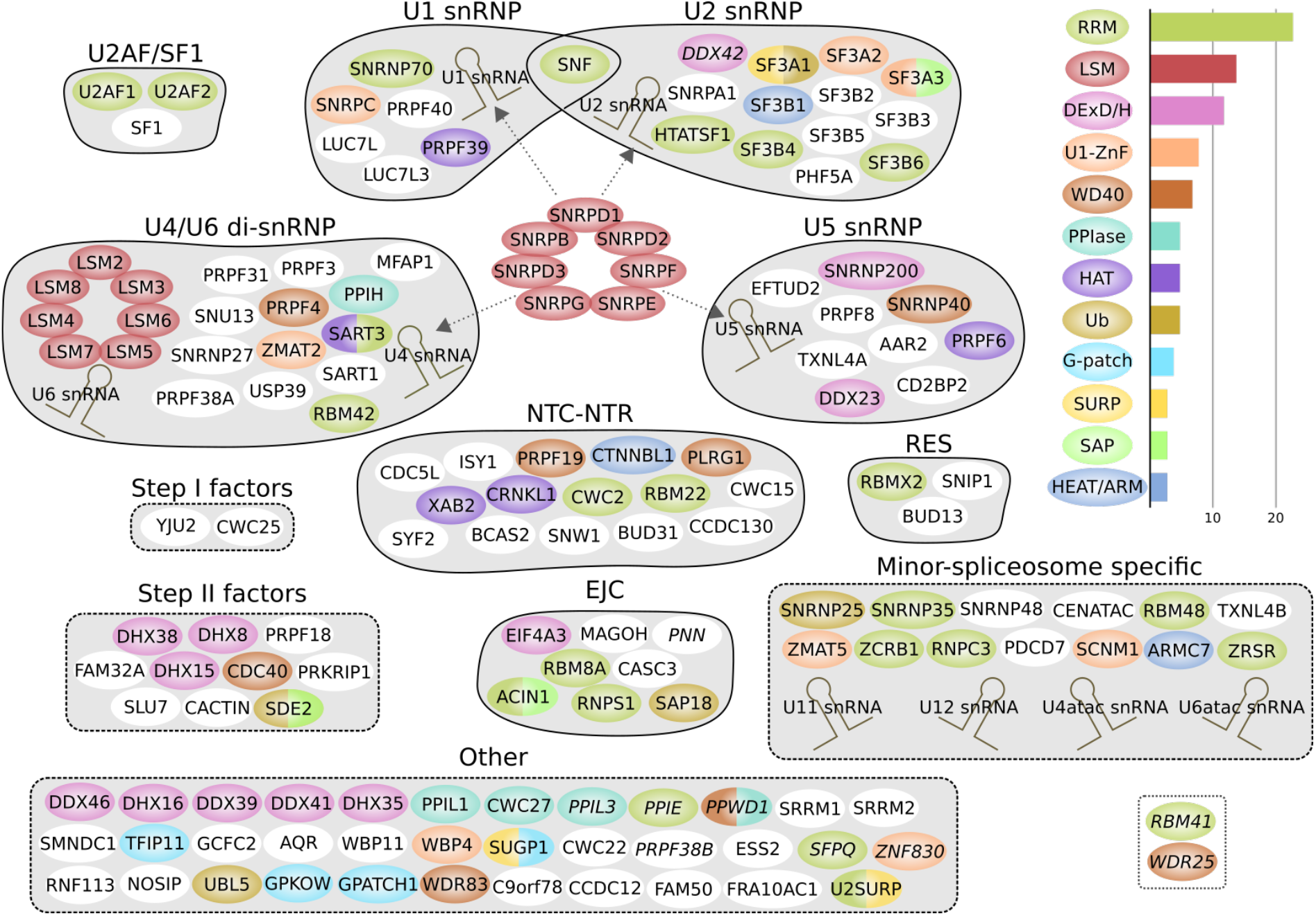
The spliceosome inferred in LECA. The names of OGs with a lower confidence score are in italics (possibly spliceosomal in LECA, see Methods). The OGs are grouped based on the subcomplex they are in or another collection (dashed line), and they are coloured based on their domain composition. Only domains that are present in at least three OGs are shown. The bar plot shows the number of OGs per domain. OGs that are only present in the minor spliceosome are displayed as minor-spliceosome specific. The main differences between the major and minor spliceosome are the presence of a U11/U12 di-snRNP instead of U1 and U2 snRNPs and the replacement of U4 and U6 snRNA with U4atac and U6atac snRNA. Two candidate minor-spliceosome specific proteins that we identified in this study are shown in the dotted box. snRNP: small nuclear ribonucleoprotein; snRNA: small nuclear RNA; NTC: Prp19-associated complex; NTR: Prp19-related complex; RES: retention and splicing complex; EJC: exon-junction complex; RRM: RNA recognition motif; ZnF: zinc finger; PPIase: peptidylprolyl isomerase; HAT: half-a-tetratricopeptide repeat; Ub: ubiquitin; HEAT/ARM: HEAT or armadillo repeats.

### From intron-encoded protein to PRPF8

As described above, the U2, U5 and U6 snRNA and the U5-snRNP protein PRPF8, as well as parts of the introns, are remnants of self-splicing group II introns. This means that during eukaryogenesis a system containing only a single RNA (the intron itself) and one protein (IEP) transformed into an enormously complex spliceosome in LECA. In principle, the prokaryotic origins of this system could be inferred from the phylogenetic affinity of IEP and the spliceosomal PRPF8 protein, as the reverse transcriptase (RT)-like domain in PRPF8 is homologous to the RT domain in IEP (Dlakić and Mushegian 2011; Qu et al. 2016; Zhao and Pyle 2016). However, phylogenetic analysis of this domain is hindered by the high sequence divergence of PRPF8 and to a lesser extent its paralog telomerase, relative to prokaryotic RT domains. In our analyses, the nuclear homologs of IEP were not clearly associated with a particular IEP type and their exact phylogenetic position in the IEP tree was unresolved (Supplementary Figure 1a).

Group II introns occur predominantly in bacteria. A recent study showed that most complete archaeal genomes do not contain group II introns, with the exception of Methanomicrobia (Miura et al. 2022). We detected group II introns in several Asgard archaeal genomes, which were from multiple different IEP types (Supplementary Figure 1b). This finding expands the set of observed IEP types in archaea to also include ML, D, E, CL2A and a separate CL type. The presence of these “bacterial” mobile elements in Asgard archaea is in good agreement with the diverse mobile elements that were recently found in circular *Heimdallarchaeum* genomes and the proposed continuous influx of bacterial genes in Asgard archaea (Wu et al. 2022). This so far unappreciated wide diversity of self-splicing group II introns in Asgard archaea might indicate the presence of such elements in the archaeal ancestor of eukaryotes.

### Expansion of the emerging spliceosome through extensive gene duplication

All other 144 spliceosomal OGs do not have a homolog in group II introns. We performed phylogenetic analyses to infer their respective evolutionary origins (Supplementary Table 4). A few OGs had a complex evolutionary history since they contain multiple domains with a separate history and resulted from a fusion event (Supplementary Information). 56 OGs were most closely related to another spliceosomal OG (Figure 2a) and therefore their pre-duplication ancestor was probably already part of the spliceosome. By collapsing such close paralogous clades of spliceosomal OGs we identified 102 ancestral spliceosomal units (Supplementary Figure 2). Duplications of spliceosomal genes increased the number of spliceosomal proteins with a factor of 1.4. The ancestral spliceosomal units themselves also originated in most cases from a duplication, but then from a gene with another function in the proto-eukaryotic cell. For 33 ancestral units we could not detect other homologs and these were therefore classified as proto-eukaryotic invention. One single spliceosomal OG, AAR2, was surprisingly found to be one-on-one orthologous to a gene in a limited number of prokaryotes, including Loki- and Gerdarchaeota (Supplementary Information). Over a hundred proteins seemed to have been recruited to the emerging spliceosome at different points during eukaryogenesis. Subsequent duplications of these proteins resulted in an even more complex spliceosome in LECA.

**Figure 2.**
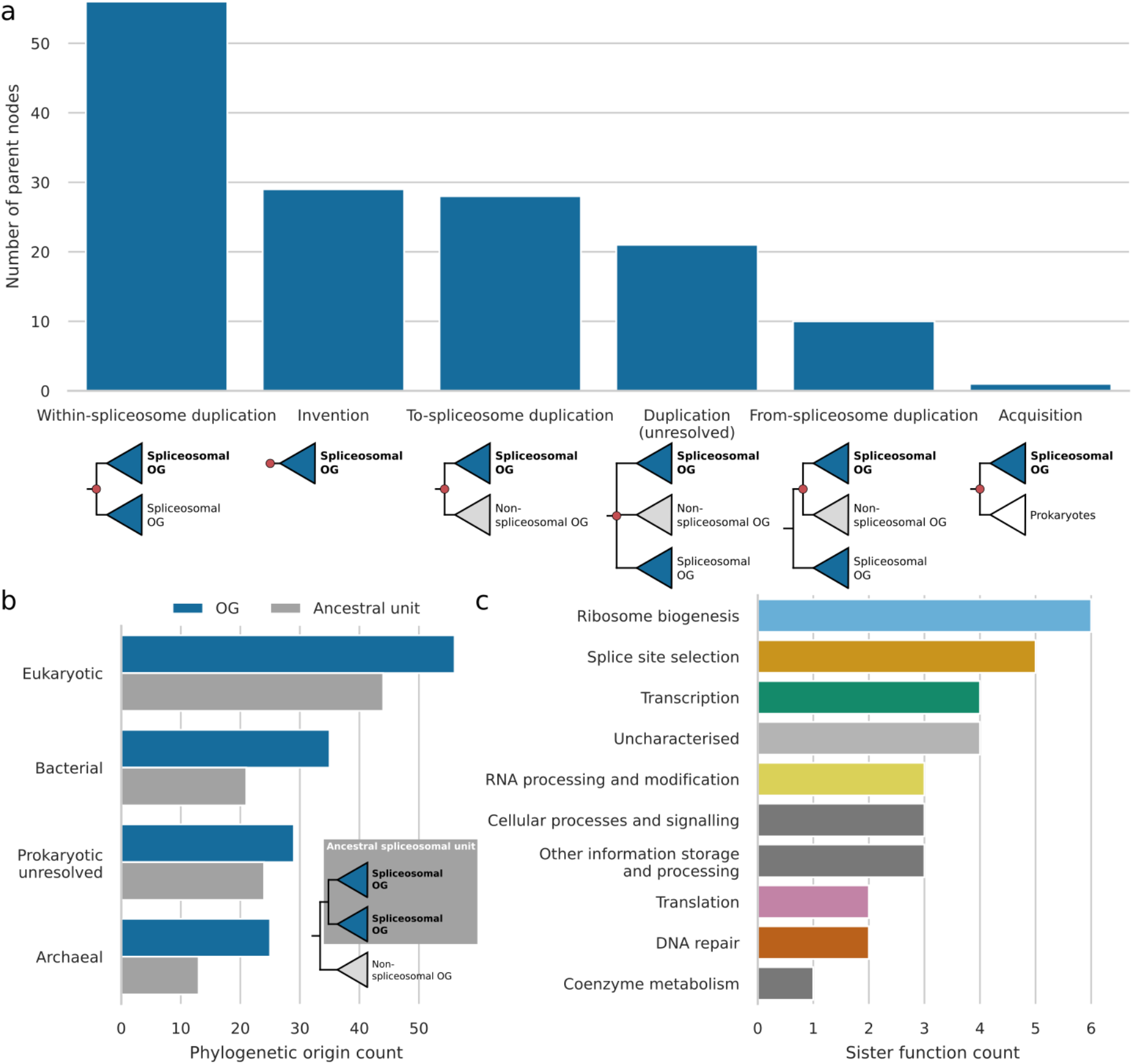
Evolutionary history of spliceosomal proteins before LECA. (a) Annotations of the parent nodes of spliceosomal OGs. These parent nodes are shown in red in the example trees below. (b) Bar plot showing the phylogenetic origins of spliceosomal OGs and ancestral spliceosomal units. (c) Functions of the sister OGs of ancestral spliceosomal units.

Eukaryotic genomes are chimeric in nature, with genes originating from the Asgard archaea-related host, the alphaproteobacteria-related protomitochondrion or other prokaryotes by means of horizontal gene transfer. The eukaryotic spliceosome mirrors this general trend. It contains considerable numbers of genes from archaeal and bacterial origin, making it a chimeric complex in phylogenetic origin (Figure 2b). The largest group, however, is comprised of genes for which we could not detect ancient homologs in prokaryotes and possibly originated *de novo*. This suggests that novel eukaryote-specific folds played a major role in shaping the emerging spliceosome. It is noteworthy that none of the acquisitions from bacteria could be traced back to alphaproteobacteria. This argues against a direct contribution of the mitochondrial endosymbiont to the spliceosome.

### Spliceosomal proteins originated predominantly from ribosomal biogenesis, translation and RNA processing proteins

A relatively large number of spliceosomal OGs were acquired from genes that functioned in ribosome biogenesis and translation (Figure 2c), especially OGs from archaeal origin. The U5 snRNP protein EFTUD2 is a paralog of elongation factor 2 (Figure 3a), which catalyses ribosomal translocation during translation elongation. The archaeal ortholog performs the same translocation function yet also probably plays a role in ribosome biogenesis that is similar to the other proto-eukaryotic paralog EFL1 (Lo Gullo et al. 2021). SNU13 and PRPF31 bind to U4 or U4atac snRNA (Nottrott et al. 2002). SNU13 is also part of the C/D-box snoRNP (Watkins et al. 2000) and PRPF31 originated from a C/D-box snoRNP protein (Figure 3b, c). The archaeal orthologs NOP5 and RPL7Ae are part of the functionally equivalent C/D box sRNP (Figure 3d), which is involved in ribosome biogenesis by modifying rRNA (Aittaleb et al. 2003; Breuer et al. 2021). The eukaryotic DDX helicases, of which six were part of LECA’s spliceosome, evolved from prokaryotic DEAD and RHLE proteins, which also function in ribosome assembly (Charollais et al. 2004; Jain 2008) (Figure 3e, f). A large group of related RNA helicases are the DHX helicases. The ancestral function of DHX helicases was probably related to ribosome biogenesis (Figure 3g). Recruitment into the spliceosome and duplications resulted in five spliceosomal DHX helicases.

**Figure 3.**
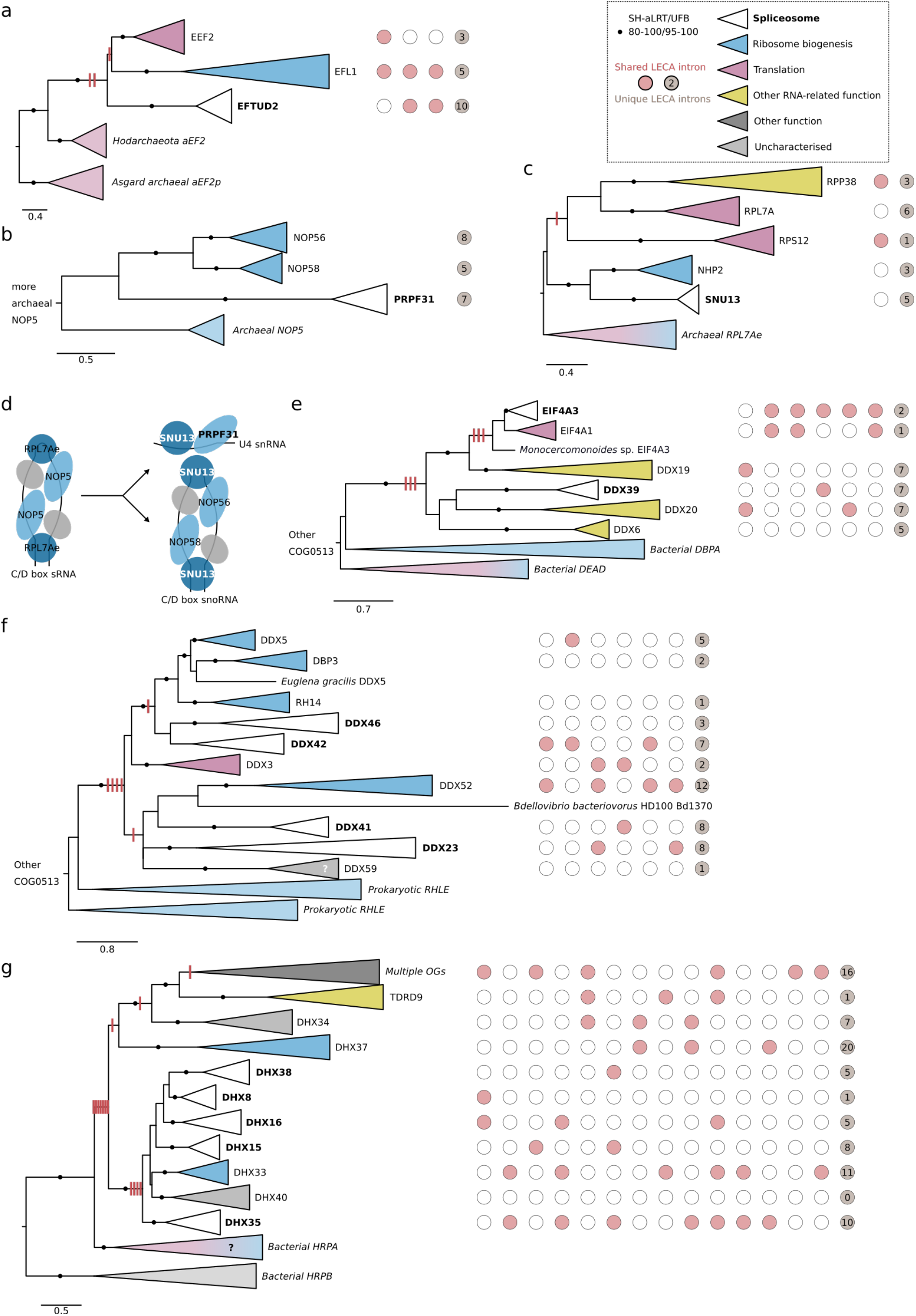
Spliceosomal proteins that originated from ribosome-related proteins. (a) Phylogenetic tree of the EF2 family. (b) Phylogenetic tree of the NOP family. (c) Phylogenetic tree of the RPL7A family. (d) Evolution of the C/D box snoRNP and U4 snRNP proteins SNU13 and PRPF31 in LECA from the C/D box sRNP in the archaeal ancestor of eukaryotes. Homologous proteins are shown in the same colour. SNU13 was present in both complexes in LECA. The grey protein corresponds with fibrillarin. (e, f) Phylogenetic tree of the DDX helicase family, displaying two separate acquisitions during eukaryogenesis in two separate panels. The function of DDX59 has not been characterised but its phylogenetic profile is similar to minor-spliceosome specific proteins (de Wolf et al. 2021) (Supplementary Figure S7b). (g) Phylogenetic tree of the DHX helicase family. (a-c, e-g) Eukaryotic LECA OGs are collapsed and coloured based on their function, as are the prokaryotic clades. Introns inferred in LECA are depicted; columns with red/white circles correspond with the presence of introns at homologous positions. The gain of introns before duplications as reconstructed using Dollo parsimony is shown with red stripes on the branches. Scale bars correspond with the number of substitutions per site. Clades with significant support as assessed with the SH-like approximate likelihood ratio (SH-aLRT) and ultrafast bootstrap (UFB) values are indicated with filled circles.

The Lsm and Sm heptamer rings that bind U6 or U6atac snRNA and other snRNAs, respectively, are also of archaeal origin. The archaeal homologs, called Sm-like archaeal proteins (SmAPs), are poorly characterised RNA-binding proteins which might function in tRNA processing and RNA degradation (Lekontseva et al. 2021). The SmAP genes are located directly adjacent to ribosomal protein RPL37e (Mura et al. 2013), emphasising the potential link with translation. The eukaryotic Lsm ring is involved in different forms of RNA processing besides splicing (Mura et al. 2013), including rRNA maturation (Kufel et al. 2003). During eukaryogenesis the Lsm ring gained the U6(atac) binding function and was recruited into the spliceosome. Subsequent gene duplications resulted in the two types of heteromeric rings of Lsm/Sm proteins in the spliceosome (Supplementary Figure 3, Supplementary Information).

A substantial fraction of the LECA spliceosome OGs contains an RNA recognition motif (RRM) (Figure 1). The proteins in this family perform diverse functions, as this domain can not only bind RNA but is also involved in protein-protein interactions (Maris et al. 2005). RRM proteins were likely acquired from a bacterium during eukaryogenesis, as proteins with this domain are present in some bacteria. Although the tree is largely unresolved due to the short length of the motif, multiple recruitments into the spliceosome can be observed, some followed by intra-spliceosome duplications (Supplementary Figure 4a). Functions of other RRM proteins that are closely related to the spliceosome OGs include transcription, splice site selection and mRNA degradation. Some OGs contain multiple RRMs, pointing at a rich history of domain and gene duplications before LECA in this family.

**Figure 4.**
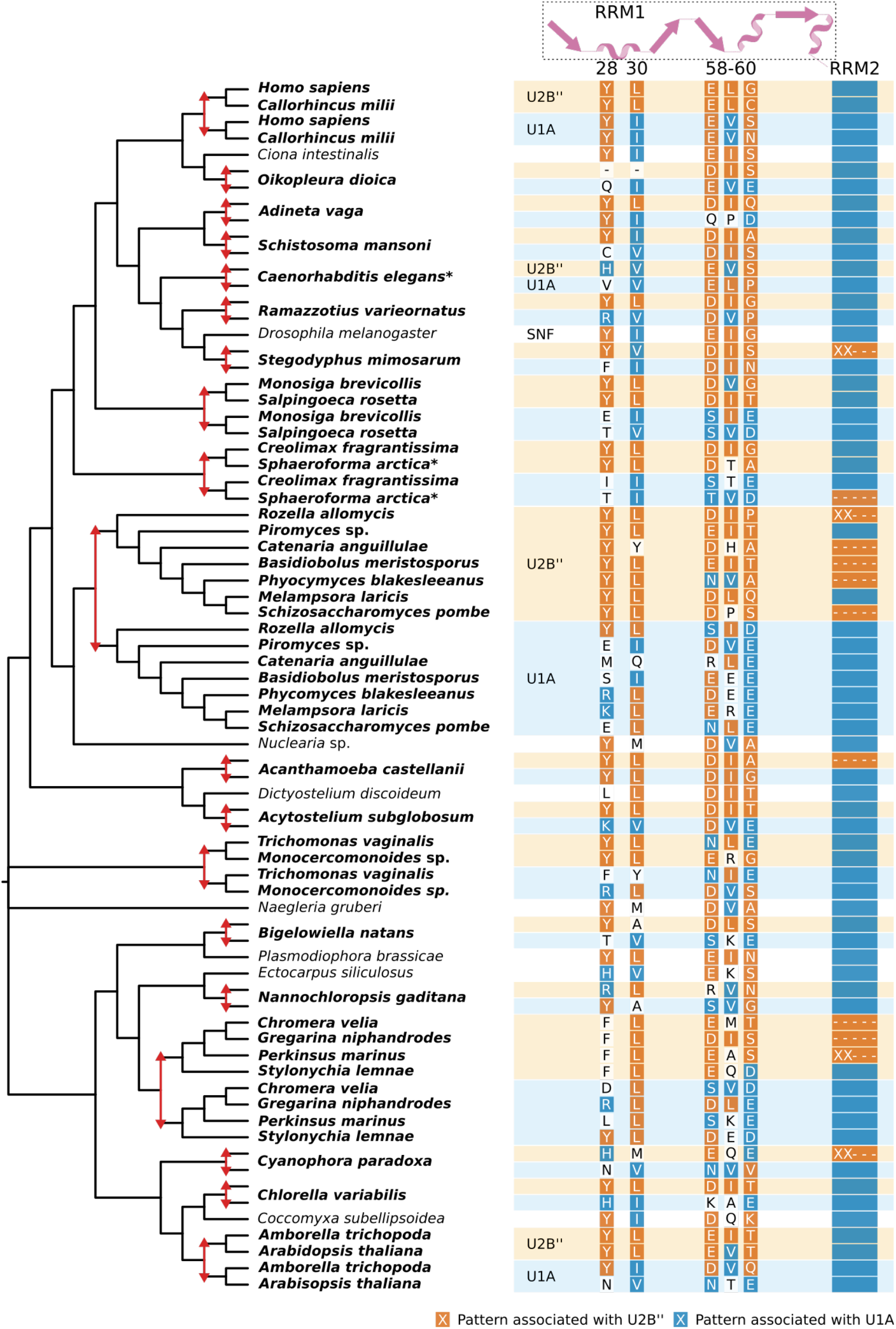
Independent gene duplications and recurrent sequence evolution in the SNF family. The reconciled tree (see Supplementary Figure 6a for the full tree) shows the positions of gene duplications (red arrows) and the species names with duplicates are in bold. The coloured rectangles next to the species names correspond with the predicted fate of the duplicates. The most prominent recurrent patterns are depicted with colours corresponding with the fate this pattern is associated with. For the second RRM (RRM2) the pattern is the presence (blue bar), absence (dashes) or partial presence (“XX---“) of this domain. The secondary structure of the first RRM (RRM1) and the position of the patterns in the D. melanogaster sequence is shown at the top. The duplications in *Sphagnum fallax* and *Emiliania huxleyi* are not shown because the duplicates are identical for the positions that are displayed.

Other large families that contributed substantially to the LECA spliceosome are the U1-type zinc finger and WD40-repeat families. The U1-type zinc finger family contains mainly spliceosomal OGs (Supplementary Figure 5a). WD40 repeats are present in many eukaryotic proteins with diverse functions. In contrast to the RNA handling functions described before, the proteins that seem to be closely related to the spliceosomal WD40 OGs are mainly involved in intracellular transport, cilia and histone modifications (Supplementary Figure 5b).

### Many minor-spliceosome specific proteins are closely related to a major spliceosome protein

The major and minor spliceosome share many subunits (Turunen et al. 2013; Bai et al. 2021) and this was very likely also the case in LECA. We inferred 13 minor-spliceosome specific proteins in LECA (Figure 1). Six of these are closely related to a major-spliceosome specific protein. The RRM proteins SNRNP35 and ZRSR have a major spliceosome equivalent as their sister paralog, SNRNP70 and U2AF1 respectively (Supplementary Figure 4b, c). RNPC3 is closely related to SNF but probably not as sister paralogs (Supplementary Figure 4d). The sister paralog of RNPC3 is RBM41, which is not well characterised. However, its phylogenetic profile corresponds with minor spliceosome OGs (Supplementary Figure 7). If RBM41 is part of the minor spliceosome, the RNPC3-RBM41 duplication would represent the only identified duplication within the minor-spliceosome specific OGs. The phylogenetic position of the other minor spliceosome OGs with an RRM is unresolved (Supplementary Figure 4a). ZMAT5 and SCNM1 are part of the U1-type zinc finger family. The equivalent of ZMAT5 in the major spliceosome is SNRPC (Will et al. 2004) and SCNM1 functions as a combination of SF3A2 and SF3A3 (Bai et al. 2021). Although the phylogenetic tree of this family is unresolved, it is likely that these major and minor spliceosome equivalents are sister paralogs. The major spliceosome OG TXNL4A and minor spliceosome OG TXNL4B are clear sister paralogs (Supplementary Figure 5c). The sister paralog of the WD40-repeat protein CDC40, called WDR25 (Supplementary Figure 5b), has a presence pattern across eukaryotes that is typical of minor spliceosome OGs (de Wolf et al. 2021), like RBM41 (Supplementary Figure 7). This protein has not been characterised either, yet its phylogenetic profile strongly suggests a function in the minor spliceosome.

A peculiar observation that we made for all major/minor pairs mentioned above is that the branch in the phylogenetic tree leading from the duplication to the minor-spliceosome specific OG is considerably shorter than the one leading to the major spliceosome-specific OG (Supplementary Figure 5d). This means that these major spliceosome-specific OGs have diverged more from the ancestral preduplication state and suggests that the function of the minor-spliceosome specific SNRNP35, ZRSR, RNPC3, ZMAT5, SCNM1, TXNL4B and possibly WDR25, better reflect the ancestral state.

### Substantial intron spread predating spliceosomal duplications

In a previous study we investigated the spread of introns in proto-eukaryotic paralogs (Vosseberg et al. 2022). Intron positions that are shared between genes that duplicated during eukaryogenesis are likely shared because they were present in the gene before it duplicated. By analysing intron positions in spliceosomal OGs we can relate the duplications in the primordial spliceosome to the spread of the elements that they function on, the introns. We therefore applied the same approach as in our previous study to the paralogs in the spliceosome. 45% of duplications that probably resulted in a novel spliceosomal gene had at least one intron traced back to the preduplication state (13 of the 29 to-spliceosome duplications). For 46% of the within-spliceosome duplications we detected shared introns between paralogs in the spliceosome (18 out of 39).

The presence of introns in ancestral genes that themselves likely did not function in the spliceosome is strikingly illustrated by the DDX and DHX helicases, with three to seven introns traced back to before the first duplication after the acquisition from prokaryotes (Figure 3). Introns shared between spliceosomal paralogs were also found in the LSM, PPIase and WD40 families (Supplementary Figure 3a, Supplementary Table 5). The U5 snRNP proteins SNRNP200 and EFTUD2, which interact with PRPF8, shared multiple introns with paralogs outside the spliceosome and likely contained introns before they became part of the spliceosome (Figure 3, Supplementary Table 5). These numbers and cases suggest that introns were already present in a substantial number of ancestral genes before the corresponding proteins were recruited into the spliceosome and subsequently duplicated within the spliceosome.

### Duplication and subfunctionalisation completed multiple times after eukaryogenesis: U1A/U2B’’

A notable difference between the LECA spliceosome and the human and yeast spliceosome is the presence of two proteins in both human and yeast stemming from a single SNF protein in LECA. In early studies the single SNF protein in *Drosophila melanogaster* was seen as the derived state and two separate proteins, U1A and U2B’’, were proposed to represent the ancestral state (Polycarpou-Schwarz et al. 1996; Williams and Hall 2010). However, with the availability of more genomes the human and yeast proteins were shown with high confidence to be the result of separate gene duplications (Williams et al. 2013). Additional SNF duplications were identified in other animal lineages (Williams et al. 2013). We observed even more independent SNF duplications, 22 in total using our set of eukaryotic genomes (Supplementary Figure 6a). *Guillardia theta* even had an additional third one, probably from the secondary endosymbiont (Supplementary Information).

*Drosophila* SNF has a dual role in the spliceosome. It is both part of the U1 snRNP, where it binds U1 snRNA, and part of the U2 snRNP, where it binds U2 snRNA and U2A’ (Weber et al. 2018). In human and yeast, U1A and U2B’’ have subfunctionalised and perform the respective functions as indicated by the snRNP in their name. To assess whether a similar subfunctionalisation has occurred in other lineages where SNF had duplicated, we looked for patterns of recurrent sequence evolution in the different paralogs with our previously published pipeline (von der Dunk and Snel 2020). Two fates could be distinguished, which we refer to as U1A and U2B’’ based on the fates in model organisms. This distinction was based on a diffuse, mainly U1A-specific signal. Upon inspection of the two fate clusters and comparison with single SNF orthologs, the fate separation seemed to be predominantly based on recurrent substitutions in the first RRM of U1A and the recurrent loss of the second RRM in U2B’’ (Figure 4). We inferred 16 RRM loss events in U2B’’-fate proteins (Supplementary Figure 6b). These recurrent sequence changes allow us to predict which inparalog is likely to have a U1A function and which one has a U2B” function in organisms where detailed biochemical studies are lacking. Besides these remarkable findings on recurrent sequence evolution, the repeated post-LECA duplications suggest that the complexification of the spliceosome by duplication during eukaryogenesis could in part have been driven by the same process as happened multiple times after LECA.

## Discussion

### A chimeric complex spliceosome that postdates the proliferation of introns

The spliceosome is one of the most complex molecular machines in present-day eukaryotes. In this study we reconstructed the composition of the spliceosome in LECA and traced the sometimes byzantine evolutionary histories of these 145 inferred spliceosomal proteins prior to LECA. Previous work has established that the core of the spliceosome - the U2, U5 and U6 snRNAs and PRPF8 – as well as the spliceosomal introns themselves evolved from self-splicing group II introns (Zimmerly and Semper 2015). Proteins of archaeal and bacterial origin were added to this core, especially proteins that performed a function in ribosome biogenesis or translation. For many proteins we could not detect other homologous proteins, suggesting that the primordial spliceosome expanded with spliceosome-specific folds. Subsequent expansions resulted from the numerous gene duplications that we observed. These duplications enabled us to assess the extent of intron positions that were shared between paralogs and likely predated the duplication event (Vosseberg et al. 2022). Our ancestral intron position reconstructions support the presence of introns in almost half of the proteins before their recruitment into the spliceosome. This suggests that introns were already widespread through the genome when most components of the complex spliceosome emerged. The increase in spliceosomal complexity did not coincide with the increase in intron numbers but followed it instead. We propose a scenario in which intragenic introns emerged early in eukaryogenesis and the complex spliceosome relatively late.

### From group II introns to a complex spliceosome

The group II introns that gave rise to the spliceosomal introns are commonly proposed to have come from the protomitochondrion (Cavalier-Smith 1991; Martin and Koonin 2006). Group II introns are present in several mitochondrial and plastid genomes (Zimmerly et al. 2001). Notwithstanding the extent of horizontal gene transfer of these introns among eukaryotes, group II introns were probably present in the mitochondria in LECA (Kim et al. 2022). Our analysis did not yield sufficient phylogenetic signal to confidently position PRPF8 in the IEP tree. However, the identification of multiple intron types in Asgard archaea makes an alternative scenario in which group II introns were present in the archaeal genome before the mitochondrial endosymbiosis also plausible (Vosseberg and Snel 2017; Vosseberg et al. 2022).

Some self-splicing introns acquired the capacity to aid the splicing of other introns. Fragments of these introns evolved into the *trans*-acting snRNAs U2/U12, U5 and U6/U6atac. IEP became a general maturase and lost its RT activity. Degeneration of self-splicing introns resulted in primordial spliceosomal introns that required these snRNAs and the general PRPF8 maturase for splicing.

Proteins involved in the assembly and functioning of another large ribonucleoprotein in the cell, the ribosome, became part of the primordial spliceosome, supplemented with other RNA-binding proteins. The evolutionary link with the ribosome emphasises the comparable composition as a ribonucleoprotein with catalysing RNA molecules (ribozymes). In contrast with the aforementioned spliceosomal snRNAs, the U1/U11 snRNA and U4/U4atac did probably not originate from the introns themselves. However, an evolutionary link with translation and rRNA processing is present for these snRNAs too. U1/U11 snRNA likely evolved from a tRNA (Hogeweg and Konings 1985). The evolutionary histories of SNU13 and PRPF31 and similarities between U4 and C/D-box RNAs suggest that the U4(atac) snRNP evolved from a C/D-box snoRNP (Watkins et al. 2000).

The contribution of gene duplications in shaping the LECA spliceosome is in line with the central role of duplications in establishing eukaryotic features during eukaryogenesis (Makarova et al. 2005; Vosseberg et al. 2021). Gene duplications were key for the emergence of spliceosome-specific proteins from proteins that were part of other complexes as well as for expanding proteins that were already part of the spliceosome. This pattern has also been observed for the kinetochore (Tromer et al. 2019). These kinetochore proteins, however, came from a wider variety of cellular processes compared with the spliceosome. The origin of another eukaryote-specific complex, the nuclear pore, compares well with the spliceosome regarding the chimeric prokaryotic ancestry of its components (Mans et al. 2004). This is unlike complexes and processes that predated eukaryogenesis, such as transcription and translation, which have a more consistent phylogenetic signal (Pittis and Gabaldón 2016; Vosseberg et al. 2021).

### Origin of two types of introns and two types of spliceosomes

Two types of introns were present in the LECA genome, U2 and U12, which were removed from the primary transcripts by the LECA major and minor spliceosome, respectively (Russell et al. 2006). The far majority of introns were probably of U2-type (Vosseberg et al. 2022). Different scenarios have been postulated for the emergence of two types of introns (Burge et al. 1998). In some scenarios the different intron types diverged from an ancestral set of introns, either in the same proto-eukaryotic lineage or two separate lineages that later fused. An alternative scenario proposes that the two types of introns originated from two separate introductions of group II introns in the genome. Previously, we called the separate introductions scenario unlikely based on the observed U12-type introns that are shared between proto-eukaryotic paralogs (Vosseberg et al. 2022). The enormous overlap in composition between the major and minor spliceosome (Turunen et al. 2013; Bai et al. 2021) refutes separate origins of these complexes from different group II introns. Many minor-spliceosome specific proteins have a close homolog in the major spliceosome and all snRNAs but U5 have equivalents in the other spliceosome type. This suggests that the divergence between the major and minor spliceosome occurred relatively late in pre-LECA spliceosome evolution, after the addition of U1 and U4 snRNA and U1 and U2 snRNP proteins. The minor-spliceosome specific proteins were estimated to have accumulated fewer substitutions after the duplications that separated major- and minor-spliceosome specific OGs. This suggests that the latter better reflect the ancestral situation. The U12-type introns and the minor spliceosome might therefore have originated earlier than the abundant U2-type introns and the major spliceosome.

### Evolution of spliceosomal complexity

During eukaryogenesis the recruitment of proteins and gene duplications resulted in an increase in spliceosomal complexity. Spliceosomal evolution after LECA is in most eukaryotic lineages dominated by simplification. A clear example is the minor spliceosome, which was lost recurrently at least 23 times (Supplementary Information). Certain lineages have experienced substantial loss of spliceosomal genes that were part of the LECA spliceosome. Only 59% of the LECA OGs are present in *Saccharomyces cerevisiae*, for example. Reduced spliceosomes have also been described in red algae and diplomonads (Hudson et al. 2019; Wong et al. 2022).

The most prominent example of a more complex spliceosome after LECA is the duplication of SNF in at least 22 lineages. To the best of our knowledge, this is the highest number of independent gene duplications in eukaryotes reported so far. It is slightly more than the 16 MadBub duplications (Tromer et al. 2016) and the 20 EF1β/δ duplications that were described before (von der Dunk and Snel 2020). We detected patterns of recurrent sequence evolution in the different paralogs, pointing at similar fates of these paralogs across eukaryotes. Given the described fates of the SNF paralogs in vertebrates, fungi, plants and *Caenorhabditis elegans*, a similar subfunctionalisation into dedicated U1 and U2 snRNP proteins in other lineages with duplications is likely.

The recurrent loss of the second RRM in proteins with a predicted U2B’’ fate suggests that the function of this RRM is mainly restricted to the U1A role. Whereas the function of the first RRM has been described as binding to U1 and U2 snRNA, the function of the second RRM has remained elusive (Williams et al. 2013). The observation of recurrent loss of this RRM in specifically U2B’’ proteins provides possible directions for further molecular research.

The dual-function SNF protein seems to be poised for duplication and subsequent subdivision of the roles in the U1 and U2 snRNP. It is tempting to speculate that the recurrent duplication of SNF indicates that this specific gene duplication and subsequent subfunctionalisation could in principle have occurred during eukaryogenesis instead. Because it did not happen to be duplicated then, it could be seen as “unfinished business” during eukaryogenesis. The cases of independent gene duplications after LECA might be used as a model for proto-eukaryotic gene duplications. Because these duplications happened relatively recent, experiments based on ancestral protein reconstructions can be performed more reliably, as has been done for the SNF family in deuterostomes (Williams et al. 2013; Delaney et al. 2014). These experiments can provide insight into the role of adaptive or neutral evolution (Finnigan et al. 2012) in creating the complex spliceosome (Vosseberg and Snel 2017).

### Investigating the emergence of the complex eukaryotic cell

Our study provides a comprehensive view on the origin of the numerous proteins in this complex molecular machine, also in relation to the spread of the introns it functions on. Further studies on the spliceosome composition in diverse eukaryotes have the potential to identify more spliceosomal proteins in LECA. New developments in detecting deep homologies (Jumper et al. 2021; Monzon et al. 2022) could reveal additional links for the spliceosomal proteins that we classified as inventions in this study. Phylogenetic analyses combined with intron analyses on the numerous other complexes that emerged during eukaryogenesis could further illuminate their origin and thereby the major transition from prokaryotes to eukaryotes.

## Materials and methods

### Data

We used a diverse set of 209 eukaryotic and 3,466 prokaryotic (predicted) proteomes, as compiled for a previous study (Vosseberg et al. 2021) from different sources (Huerta-Cepas et al. 2016; Zaremba-Niedzwiedzka et al. 2017; Deutekom et al. 2019). Proteins from 167 of the eukaryotic species had been grouped in OGs using different approaches (Deutekom et al. 2021). To illuminate the evolutionary history of some protein families (see below) we made use of the widely expanded set of Asgard archaeal genomes that has come available since. By including genomes from numerous studies (Liu et al. 2018; Tully et al. 2018; Huang et al. 2019; Seitz et al. 2019; Imachi et al. 2020; Farag et al. 2021; Liu et al. 2021; Sun et al. 2021; Zhao and Biddle 2021; Wu et al. 2022), the number of Asgard archaeal proteomes in our expanded set amounted to 133 in total. If no predicted proteome was available, the genomes were annotated with Prokka v1.13 (Seemann 2014) for the genomes from (Liu et al. 2018; Seitz et al. 2019) or v1.14.6 with the metagenome option for the genomes from (Farag et al. 2021; Liu et al. 2021).

### Reconstructing LECA’s spliceosome

To infer the composition of the spliceosome in LECA, we searched for orthologs of proteins in the well-studied *Homo sapiens* and *Saccharomyces cerevisiae* spliceosome complexes in other eukaryotic proteomes. A list of human and budding yeast spliceosomal proteins was obtained from the UniProt database (The UniProt Consortium 2019) on 26 February 2020, only including manually reviewed proteins (Supplementary Table 1, 2). Proteins that are involved in other processes (such as transcription and polyadenylation) and splice site selection and splicing regulation were removed. The list was supplemented with human spliceosomal proteins from recent literature (Bai et al. 2021; Sales-Lee et al. 2021; de Wolf et al. 2021). Initial evolutionary scenarios of these proteins were inferred based on the approach of Van Hooff *et al*. (2019) (van Hooff et al. 2019). In short, the human and yeast protein sequences were searched against our in-house eukaryotic proteome database (Deutekom et al. 2019) with blastp (Altschul et al. 1990). Significant hits (E-value 0.001 or lower) in *H. sapiens, Xenopus tropicalis, D. melanogaster, Salpingoeca rosetta, S. cerevisiae, Schizosaccharomyces pombe, Spizellomyces punctatus, Thecamonas trahens, Acanthamoeba castellanii, Dictyostelium discoideum, Arabidopsis thaliana, Chlamydomonas reinhardtii, Cyanidioschyzon merolae, Ectocarpus siliculosus, Plasmodium falciparum, Plasmodiophora brassicae, Naegleria gruberi, Leishmania major, Giardia intestinalis* and *Monocercomonoides* sp. were aligned with MAFFT v7.310 (E-INS-i option) (Katoh and Standley 2013). These alignments were trimmed with trimAl v1.4.rev15 (gappyout option) (Capella-Gutiérrez et al. 2009) and a phylogenetic tree was inferred with IQ-TREE v1.6.4 (Nguyen et al. 2015) using the LG+G4 model to establish the initial scenario (van Hooff et al. 2019): (i) easy, in case of orthologs in a diverse set of eukaryotes; (ii) ancient (pre-LECA) duplication, when the set of homologs also includes clades of more distantly related homologs across eukaryotes; (iii) lineage-specific (post-LECA) duplication, when the spliceosomal function likely originated after LECA ; (iv) taxonomically limited, with homologs in a limited set of eukaryotes. The latter cases were further studied by checking hits in the complete set of eukaryotes. For SNRNP27, CASC3 and WBP11, hits to the more sensitive Pfam models PF08648, PF09405 and PF09429 (Finn et al. 2016) detected before (Vosseberg et al. 2021) were used instead of the BLAST-based homologs.

In case of an easy or ancient duplication scenario, a LECA OG was defined. If members of this OG were present in both the human and yeast spliceosome, it was classified as a LECA spliceosome OG. Yeast LIN1 (CD2BP2 ortholog) and PRP24 (SART3 ortholog) and human LUC7L and LUC7L2 (LUC7 orthologs) were not in the initial set but their ortholog was. These were included in the original list because these were also clearly described as spliceosomal in the literature. If an ortholog was not present in yeast, spliceosomal annotations for orthologs in *S. pombe, A. thaliana* (both in the UniProt database) or *Cryptococcus neoformans* (Sales-Lee et al. 2021) were checked. If an ortholog was not present in human, the function of the *A. thaliana* ortholog was investigated. If these orthologs were not characterized, they were classified as spliceosomal in LECA if their close paralog was also in the spliceosome, or if they only had an annotated spliceosomal function. If their main function was in the spliceosome or if they were not well-characterised, they were classified as possibly spliceosomal. In case of multiple functions, the OG was discarded. The reconstruction of spliceosome OGs in LECA is summarised in Supplementary Table 3.

### Inferring pre-LECA evolutionary histories

To trace the pre-LECA histories of the inferred spliceosomal LECA proteins we performed phylogenetic analyses of these proteins with other eukaryotic OGs and with prokaryotic proteins that are homologous to the spliceosomal proteins. We started by analysing the domain composition of the proteins and looking for these domains or full-length proteins in trees that we created for a previous study (Vosseberg et al. 2021). Additional phylogenetic analyses were performed for the families described below. Multiple sequence alignments were made with MAFFT v7.310 (Katoh and Standley 2013) and subsequently trimmed to remove parts of the alignment of low quality with trimAl v1.4.rev15 (Capella-Gutiérrez et al. 2009) or Divvier v1.0 (Ali et al. 2019) (maximum of 50% gaps per position). The chosen options per family are shown in Supplementary Table 6. Phylogenetic trees were inferred using IQ-TREE v2.1.3 (Minh et al. 2020) with the best substitution model among nuclear models including LG+C{10,20,30,40,50,60} mixture models identified by ModelFinder (Kalyaanamoorthy et al. 2017). Mixtures models with an F-class were not considered, as recently recommended (Baños et al. 2022). Branch supports were calculated with 1,000 ultrafast bootstraps (Hoang et al. 2018) and the SH-like approximate likelihood ratio test (Guindon et al. 2010). Topologies were compared using the approximately unbiased test (Shimodaira 2002) with 10,000 replicates.

#### IEP-PRPF8

Representative sequences of prokaryotic and organellar IEP sequences and other prokaryotic RT-containing sequences were chosen from two datasets (Candales et al. 2012; Toro and Nisa-Martínez 2014) and supplemented with four Asgard archaeal IEP sequences (Zaremba-Niedzwiedzka et al. 2017). We also selected slowly evolving representatives for PRPF8 and TERT. For the tree that included PRPF8 and TERT separate alignments were made for the prokaryotic and organellar (E-INS-i algorithm), PRPF8 and TERT sequences (both with L-INS-i). We extracted the RT fingers-palm and thumb domains from these alignments based on a published structural alignment (Qu et al. 2016). The extracted domains were aligned and a tree was inferred. A constrained tree search with a monophyletic PRPF8 and TERT clade was additionally performed.

We used eggNOG 4.5 (Huerta-Cepas et al. 2016) annotations to identify additional Asgard archaeal IEPs by executing emapper-1.0.3 (Huerta-Cepas et al. 2017) with DIAMOND v0.8.22.84 (Buchfink et al. 2015) searches on the expanded Asgard set. Proteins assigned to COG3344 were combined with the selection of IEP sequences; non-IEP COG3344 hits were discarded based on a preliminary phylogenetic tree.

#### AAR2

Only three prokaryotic AAR2 homologs were detected in the initial dataset based on hits to the PF05282 model (Vosseberg et al. 2021), one in *Limnospira maxima* and two in *Lokiarchaeum*. We used the same approach to identify detect additional hits in the expanded set of Asgard archaea by running hmmsearch (HMMER v3.3.2 (Eddy 2011)) with the Pfam 31.0 hidden Markov models (HMMs) (Finn et al. 2016) using the gathering thresholds. Additionally, hmmsearch with the PF05282.14 model was performed on the EBI server (http://www.ebi.ac.uk/Tools/hmmer/search/hmmsearch) www.ebi.ac.uk/Tools/hmmer/search/hmmsearch) against the UniProtKB database on 21 April 2022.

#### EFTUD2

The EF2 family has underwent multiple duplications in archaeal and eukaryotic evolution resulting in two orthologs in the last Asgard archaeal common ancestor and three in LECA (Narrowe et al. 2018). The latter are represented in eukaryotic eggNOG families (euNOGs) KOG0467, KOG0468 and KOG0469. To increase the phylogenetic resolution we used a ScrollSaw-inspired approach (Elias et al. 2012; van Wijk and Snel 2020; Vosseberg et al. 2021) to select slowly evolving sequences from four main eukaryotic clades (Amorphea, Diaphoretickes, Discoba and Metamonada). Asgard archaeal sequences assigned to COG0480 were aligned with E-INS-i. The alignment was trimmed with trimAl (-gt 0.5) and a tree was inferred using the LG+G4 model. Hodarchaeal representatives and other Asgard sequences from the same Asgard archaeal OG (see Supplementary Information) were combined with the eukaryotic sequences.

#### PRPF31 and SNU13

For PRPF31, the sequences in the PF01798 tree were replaced with the corresponding full-length sequences to increase the phylogenetic signal. Based on the PF01248 tree, which includes SNU13, we chose two slowly evolving Opimoda and two Diphoda sequences (Derelle et al. 2015) per OG, supplemented with the archaeal RPL7Ae sequences. Full-length sequences were used for subsequent phylogenetic inference.

#### DDX helicases

Slowly evolving eukaryotic DDX helicase sequences were selected using the ScrollSaw-based approach on the sequences that were assigned to euNOGs that were part of the COG0513 cluster (Makarova et al. 2005). An alignment of these sequences was created (E-INS-i, trimAl -gt 0.5) and a phylogenetic tree was inferred with FastTree v2.1.10 (LG model) (Price et al. 2010). From this tree we selected per OG the sequence on the shortest branch for each of the four eukaryotic clades (if present and not on a deviating long branch). The selected sequences were split into the two acquisitions and combined with prokaryotic COG0513 representatives.

#### DHX helicases

A similar approach as for the DDX helicases was applied to the COG1643 cluster (Makarova et al. 2005). The initial tree was based on an alignment created with E-INS-i and trimAl (gappyout option) and made using the LG+F+R8 model in IQ-TREE. An unclear clade with multiple OGs was reduced and sequences from the missing DHX40 OG were added.

#### LSM

To elucidate the pre-LECA history of the Lsm/Sm proteins we initially made a tree combining the eukaryotic sequences from LECA OGs in the Sm-like Pfam clan (PF01423, PF12701 and PF14438). We selected slowly evolving sequences as described for the DDX and DHX helicases from the resulting tree (alignment with FFT-NS-I, trimming with trimAl (-gt 0.1), tree with the LG+G4 model). LSM14 and ATXN2 were not included in the selection because of their divergent nature. The full-length sequences in the expanded set of Asgard archaea that were PF01423 hits were used for the SmAP tree. We selected representatives from the different clades and combined these with the full-length versions of the previously selected eukaryotic sequences. We also performed a constrained tree search with one monophyletic eukaryotic clade.

#### RRM and TXNL4

We identified LECA OGs in the PF00076 (RRM) tree based on automatic annotation and manual assessment (i.e., a high support value and substantial pre-LECA branch length). Per OG the shortest Opimoda and Diphoda sequence on the shortest branch were selected. For the different subtrees we selected full-length sequences in the OGs from *H. sapiens, A. castellanii, A. thaliana, Aphanomyces astaci, Monocercomonoides* sp. and *N. gruberi*. For RBM41 the *Selaginella moellendorffii* sequence was included to replace the missing *A. thaliana* ortholog. To illustrate the relationship between TXNL4A and TXNL4B in the larger thioredoxin family, we used orthologs from the same species as chosen for the RRM subtrees.

#### U1-type zinc finger

Slowly evolving sequences from the euNOGs in the smart00451 cluster (Makarova et al. 2005), supplemented with the SCNM1 euNOG ENOG410IW6J, were selected with the aforementioned ScrollSaw-based approach. These sequences were aligned with the E-INS-i algorithm and the resulting alignment was trimmed with trimAl (-gt 0.25). Based on the inferred tree with the VT+R4 model, we selected the shortest sequences per OG from each of the four eukaryotic groups.

#### WD40

The ScrollSaw-based approach was also applied to the euNOGs in the COG2319 cluster (Makarova et al. 2005), using bidirectional best hits between Opimoda and Diphoda species instead because of the size of this protein family. An alignment of the selected sequences was made (E-INS-i, trimAl gappyout) and a tree inferred (LG+R4 model). Per OG the shortest Opimoda and Diphoda sequence was chosen. PPWD1 and some potential sister OGs based on the BLAST trees were not in the COG2319 cluster. We followed a similar approach to identify slowly evolving sequences for these euNOGs (KOG0882, ENOG410IQTX, -0KD7K and -0IF90), using a different gap threshold (50%) and substitution model (LG+R3). Based on the BLAST trees and the COG2319 cluster tree, we identified potential sister OGs and inferred a tree with these OGs and the spliceosomal OGs.

### Ancestral intron position reconstructions

We performed ancestral intron position reconstructions for the identified pre-LECA paralogs in the entire clade or only for the spliceosomal OGs and sister OGs (Supplementary Table 5), depending on the number of OGs in an acquisition or invention. To establish the content of the OGs, we started with the euNOG assignments. If the taxonomic distribution of the euNOG was limited, we continued with the Broccoli (Derelle et al. 2020) OG assignments (Deutekom et al., 2021). A phylogenetic tree of the OG was inferred to check for the presence of non-orthologous or dubious sequences and remove these (E-INS-i, trimAl -gt 0.5 or -gappyout, FastTree -lg). After cleaning up the OGs, a final E-INS-i alignment was made. Except for the alignment with PRPF8 and TERT, which was based on the RT domain (see ‘IEP-PRPF8’ above), the full-length sequences were used for this alignment. Intron positions were mapped onto the alignment using the method described before (Vosseberg et al. 2022). LECA introns were inferred with Malin (Csűrös 2008) using the intron gain and loss rates that we previously estimated for the KOG clusters (Vosseberg et al. 2022). Pre-duplication introns were inferred using Dollo parsimony.

### Recurrent duplication and subfunctionalisation of SNF

To identify post-LECA duplications, SNF sequences were aligned with E-INS-i and this alignment was trimmed with Divvier. The SNF tree was inferred with the LG+C50+R6 model and manually reconciled with the species tree to annotate gene duplication events. We looked at potential duplications in more detail by remaking trees of specific parts of the tree, including additional species from our original set (Deutekom et al. 2019). Prior to making the final alignment, we removed additional in-paralogs, probable fission events or partial annotations and the sequences from *Guillardia theta*, which had likely acquired a third copy from its endosymbiont. The final alignment was made with the E-INS-i algorithm. This alignment and the annotated duplication events were used as input for our previously published pipeline to identify patterns of recurrent sequence evolution after independent gene duplications (von der Dunk and Snel 2020).

### Statistical analysis

Statistical analyses were performed in Python using NumPy v1.21.141 (Harris et al. 2020) and pandas v1.3.142 (McKinney 2010). Figures were created with Matplotlib v3.4.245 (Hunter 2007), seaborn v0.11.146 (Waskom 2021) and FigTree v1.4.3 (https://github.com/rambaut/figtree).

### Data availability

Fasta files, phylogenetic trees and mapped intron files are available in figshare (https://doi.org/10.6084/m9.figshare.20653575).

## Acknowledgements

We thank the members of the Theoretical Biology & Bioinformatics group for useful discussions. This work is part of the research programme VICI with project number 016.160.638, which is financed by the Netherlands Organisation for Scientific Research (NWO).

## Author contributions

J.V. and B.S. conceived the study. J.V. and D.S. performed the research. S.H.A.v.d.D. aided with the recurrent sequence evolution analysis of the SNF family. J.V., D.S. and B.S. analysed and interpreted the results. J.V. wrote the manuscript, which was edited and approved by the other authors.

## Notes

### Competing Interest Statement

The authors have declared no competing interest.

