## Supplementary Information for "Integrating phylogenetics with intron positions illuminates the origin of the complex spliceosome"

### Supplementary Results and Discussion

*Gene fusion*

Some spliceosomal OGs had a more complex evolutionary history because their domain composition was the result of a gene fusion during eukaryogenesis. Three OGs combine an RRM with another domain that is also part of other spliceosomal OGs: SART3, ACIN1 and U2SURP (Figure 1). The statistics in the main text (e.g., sister function and phylogenetic origin) are based on the largest domain, namely the HAT domain for SART3 and the RRM domain for ACIN1 and U2SURP, instead of SAP and SURP, respectively. These SAP and SURP domains are present only in combination with other domains in the spliceosomal OGs. Ubiquitin domains are combined with a SURP domain in SF3A1 and with a SAP domain in SDE2. The main classification for both was based on the ubiquitin domain. In SF3A3, a SAP domain is combined with a U1-type zinc finger and a SURP and G-patch domain are combined in SUGP1. For these proteins, the SURP and SAP domain were respectively used for the statistics. The G-patch domain was also not used for this in case of TFIP11, which combines this domain with a GCFC domain. PPWD1 combines a WD40 and PPIase domain, with the former comprising a larger part of the protein. SNRNP200 has an additional PWI domain. PRPF8 acquired a MPN domain, which is present in proteins involved in deubiquitination, from an Asgard archaea-derived protein.

A specific example of a domain that was acquired after LECA and is worth mentioning is the PPIase domain in PPIE, which was acquired in the opisthokont lineage. The OG name PPIE is due to the PPIase domain in the human protein, despite only the RRM domain probably being present in LECA.

*AAR2*

The presence of 1-on-1 orthologs of eukaryotic AAR2 in prokaryotes is remarkable since the spliceosome is eukaryote-specific. AAR2 binds to parts of PRPF8 that are not present in IEP, including the RNaseH-like domain (Galej et al. 2013). It has a very sparse presence distribution in prokaryotes. We detected homologs in multiple cyanobacteria, Lokiarchaeota, Gerdarchaeota, one Helarchaeote, one unclassified Asgard archaeon, one planctomycete and one bacterium classified as Ardenticatenales (Chloroflexi). These proteins have not been characterised yet. Their function could provide insight into the transition from AAR2's original prokaryotic function to its spliceosomal function in eukaryotes.

*Asgard archaeal EF2*

The evolutionary history of EF2 in archaea involved a gene duplication before the last Asgard archaeal common ancestor (Narrowe et al. 2018). The LC3 lineage, which has recently been renamed to Hodarchaeota (Liu et al. 2021), lost one of the paralogs and concomitantly was the only Asgard archaeal lineage to retain the diphthamide biosynthesis genes (Narrowe et al. 2018). In our tree of archaeal EF2 sequences we could distinguish two clear OGs, which corresponded with the "bona fide" EF2 (aEF-2) and EF2 "paralog" (aEF-2p) described before (Narrowe et al. 2018). Interestingly, the Hodarchaeal sequences were with high support in the aEF-2p clade. The Jordarchaea also encoded solely EF2 sequences from this OG and had retained the HRG motif, which is present in archaea with a single EF2 copy. This motif is the site of diphthamide modification. In fact, the diphthamide biosynthesis COGs COG1736, -1798 and -2102 were present in Hodarchaea, Jordarchaea and the Asgard Lake Cootharaba group (ALCG). For the latter group, no EF2 homologs were detected, probably because of incomplete genomes. These findings corroborate the previously published pattern of losing one of the paralogs and retaining the diphthamide biosynthesis genes in Asgard archaea and Korarchaeota (Narrowe et al. 2018).

Eukaryotic EF2, EFL1 and EFTUD2 are strongly affiliated to the archaeal EF2 from the LC3 lineage. According to our analysis this would suggest that the aEF-2p sequences retained their ribosome translocation function and HRG motifs after duplication in at least the lineages leading to the eukaryotes, Hodarchaea, Jordarchaea and probably ALCG. In these same lineages the diphthamide biosynthesis genes were retained and the "bona fide" EF2 was lost. Alternative scenarios with multiple HGT events among Asgard archaea are less likely since it involves multiple genes.

*Lsm*

The interpretation of the Lsm tree, especially when including a wide range of archaeal sequences, is notoriously difficult due to its unresolvedness. This likely results from a low phylogenetic signal in these relatively short sequences. Our phylogenetic tree from only eukaryotic sequences is not dissimilar to the previously observed Sm/Lsm pairing of proteins with the same position in the Sm and Lsm rings (Veretnik et al. 2009) (Supplementary Figure 3a). This provides support to the co-duplication of the genes that were part of an ancestral heteroheptameric ring, resulting in separate Sm and Lsm rings. Subsequent duplication of a few single Sm and Lsm genes resulted in two other Lsm/Sm rings in LECA. The Lsm rings are involved in multiple RNA-related processes (Scofield and Lynch 2008), including U6 snRNA binding, and therefore likely represent the ancestral function of these proteins.

Asgard archaeal SmAP genes can be separated in three OGs that were probably present in the Asgard ancestor (Supplementary Figure 3b). The eukaryotic sequences were split in two groups that were related to two separate Asgard archaeal OGs (Supplementary Figure 3c). Although eukaryotic monophyly could not be rejected (approximately unbiased test, $P$ = 0.225), eukaryotic Sm genes seem to originate from two separate host genes.

*SNF*

The SNF tree is mostly unresolved and proteins that have the same predicted fate cluster together to some extent (Supplementary Figure 6a). This suggests that there is a conflicting signal between the phylogenetic signal and the fate-specific patterns. The gene duplications that occurred in the ancestors of Tracheophyta, *S. fallax*, *A. castellanii*, *E. huxleyi*, *C. elegans* and *Stegodyphus mimosarum* are clear from the tree. The duplicates in *Chlorella variabilis*, *Acytostelium subglobosum* and *Nannochloropsis gaditana* are close together in the tree, albeit not monophyletic, making these likely the result of lineage-specific duplications. The previously described duplication in vertebrates can also be seen in the SNF tree (Williams et al. 2013). The paralogs in *Oikopleura dioica*, *Adineta vaga*, *Ramazzottius varieornatus*, *Cyanophora paradoxa* and *Bigelowiella natans* are quite divergent and do not cluster together but probably represent lineage-specific dupliations. The two copies in *Schistosoma mansoni* likely resulted from an ancestral duplication in the Platyhelminthes, either before or after the split with *Schmidtea mediterranea*. Two SNF genes were present in the ancestors of Ichthyosporea, Choanoflagellatea and probably Fungi, likely reflecting separate duplications. An alternative scenario in which a gene duplication took place in an opisthokont ancestor and subsequently one of the copies was lost in the animal, *Capsaspora owczarzaki* and Nuclearia lineages is less likely, as it requires a dual U1 and U2 function to have been maintained in these two paralogs until those lineages separated from the ones that kept both copies. Given the presence of two SNF genes in the Alveolata and Metamonada species that we considered, duplications before the last common ancestors of Alveolata and Metamonada, respectively, is the most parsimonious explanation. In a tree with only Archaeplastida and Cryptista sequences one of the three *G. theta* sequences was together with the red algae, indicating an endosymbiotic origin of the third SNF sequence in this species.

Using the pipeline that we published before (von der Dunk and Snel 2020), we obtained a high pervasiveness score ($P$ = 21, including the excluded *G. theta* would make 22 independent duplication events) and low fate similarity score ($Z_F$ = 1.867). The predicted fates of vertebrate, yeast and *A. thaliana* genes were consistent with the U1A and U2B'' distinction. However, the *C. elegans* paralogs RNP-3 and RNP-2 were clustered with the U1A and U2B'' proteins, respectively, contrary to their functional characterisation (Saldi et al. 2007). This could be due to the functional redundancy observed for these paralogs (Saldi et al. 2007). We adjusted the fate prediction for the *C. elegans* proteins to fit their described function. For three species the predicted fates did not correspond with the fates of orthologs in closely related species. The U2B'' prediction of the U1A protein in *Sphaeroforma arctica* was probably caused by its missing second RRM (Figure 4, Supplementary Figure 6b). The prediction of the SNF proteins in *Coemansia reversa* and *Ichthyophthirius multifiliis* was also not consistent with close fungal and alveolate orthologs, respectively. The not fully consistent prediction is not unexpected given the low fate similarity.

Besides the recurrent substitutions in U1A and U2B'' fate proteins, lineage-specific duplications could have played a role in the subfunctionalisation. Biochemical analyses of reconstructed ancestral sequences in vertebrates demonstrated how substitutions of five consecutive amino acids in the first RRM effected the changes in snRNA specificity (Delaney et al. 2014). Recurrent substitutions in these positions were not identified by our pipeline.

*Minor spliceosome*

Based on the absence of both minor spliceosome-specific proteins and snRNAs we could infer that the minor spliceosome was lost completely 23 times, in the Ichthyophonida, Choanoflagellatea, Chromadorea, *R. varieornatus*, *O. dioica*, *Nuclearia* sp., *Mitosporidium daphniae*, Blastocladiomycota, Kickxellomycotina, *Mortierellomycotina elongata*, Dikarya, Entamoebidae, Haptophyta, Labyrinthulea, Ochrophyta, Myzozoa, Oligohymenophorea, Rhizaria, *G. theta*, Rhodophyta, Chlorophyta, Metamonada and Discoba (if these last two groups are not monophyletic). Additionally, the loss of the minor spliceosome is likely in *D. discoideum, A. subglobosum* and *T. trahens* (only RNPC3 detected in those species) and *Encephalitozoon intestinalis* (only ZMAT5 detected). The latter would make the loss in both *M. daphniae* and *E. intestinalis* likely to have happened in a microsporidian ancestor. This parsimonious reconstruction of minor spliceosome loss suggests 10 additional loss events compared with a previous study (López et al. 2008).

### Supplementary Figures

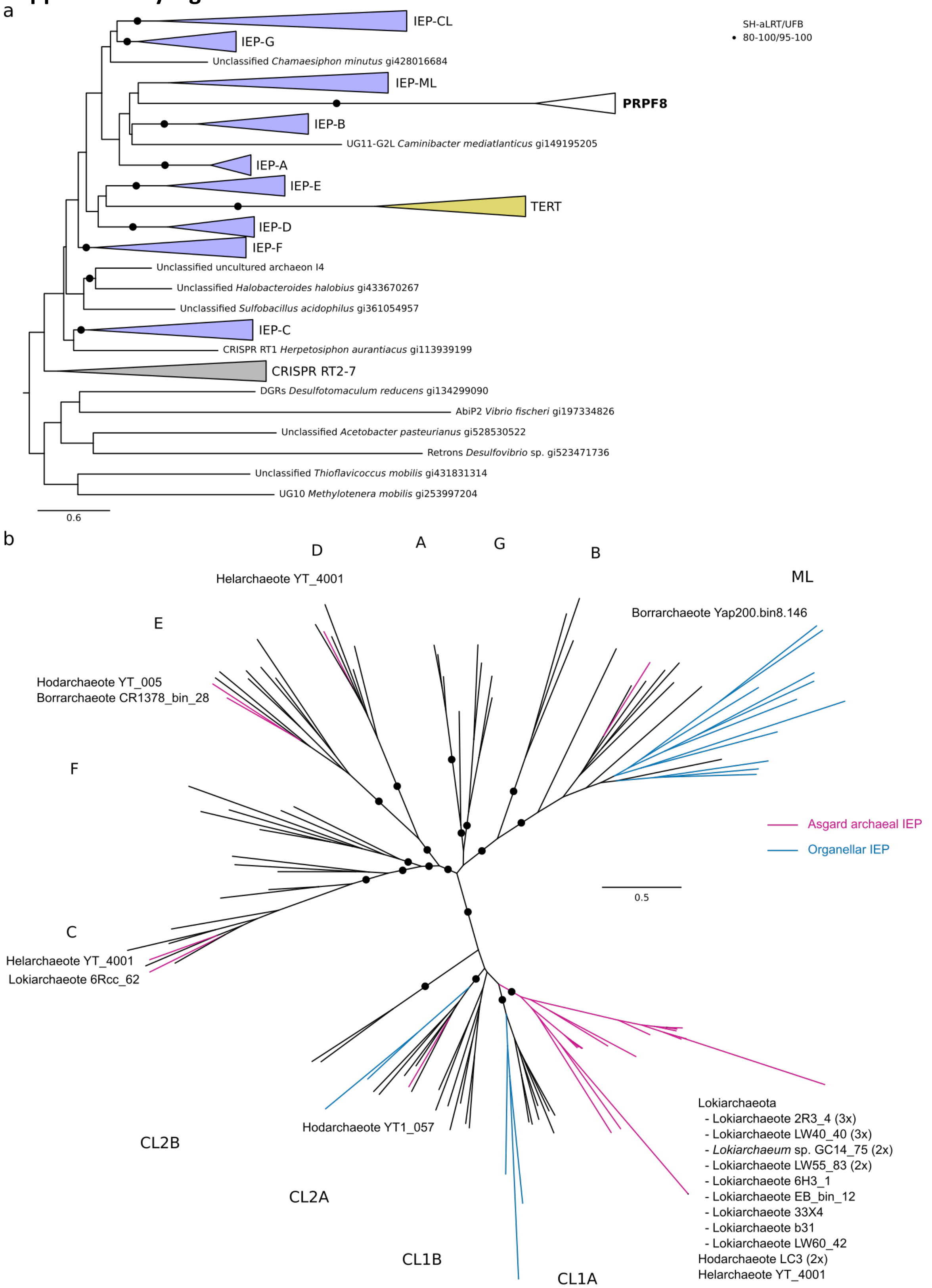


Supplementary Figure 1. Evolutionary history of the intron-encoded protein.
(a) Phylogenetic position of PRPF8 and TERT in the IEP tree. (b) Phylogenetic position of Asgard archaeal IEPs. Filled circles correspond with an SH-like approximate likelihood ratio of at least 0.8 and an ultrafast bootstrap value of at least 0.95; scale bars represent the number of substitutions per site. ML: mitochondria-like, CL: chloroplast-like, RT: reverse transcriptase, DGRs: diversity-generating retroelements, UG10 and UG11: RTs of unknown function.

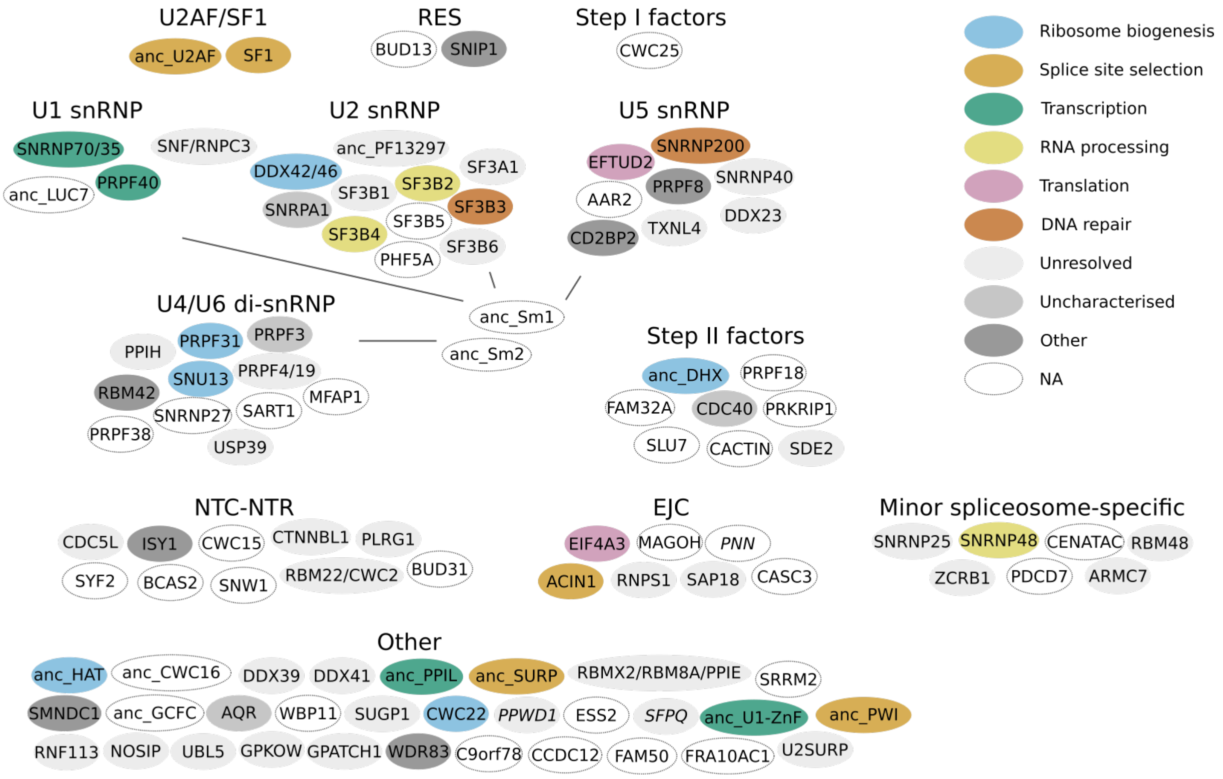


**Supplementary Figure 2. Ancestral spliceosomal units.**
Duplications within the spliceosome are collapsed and the function of the eukaryotic sister OG (if detected) is indicated.

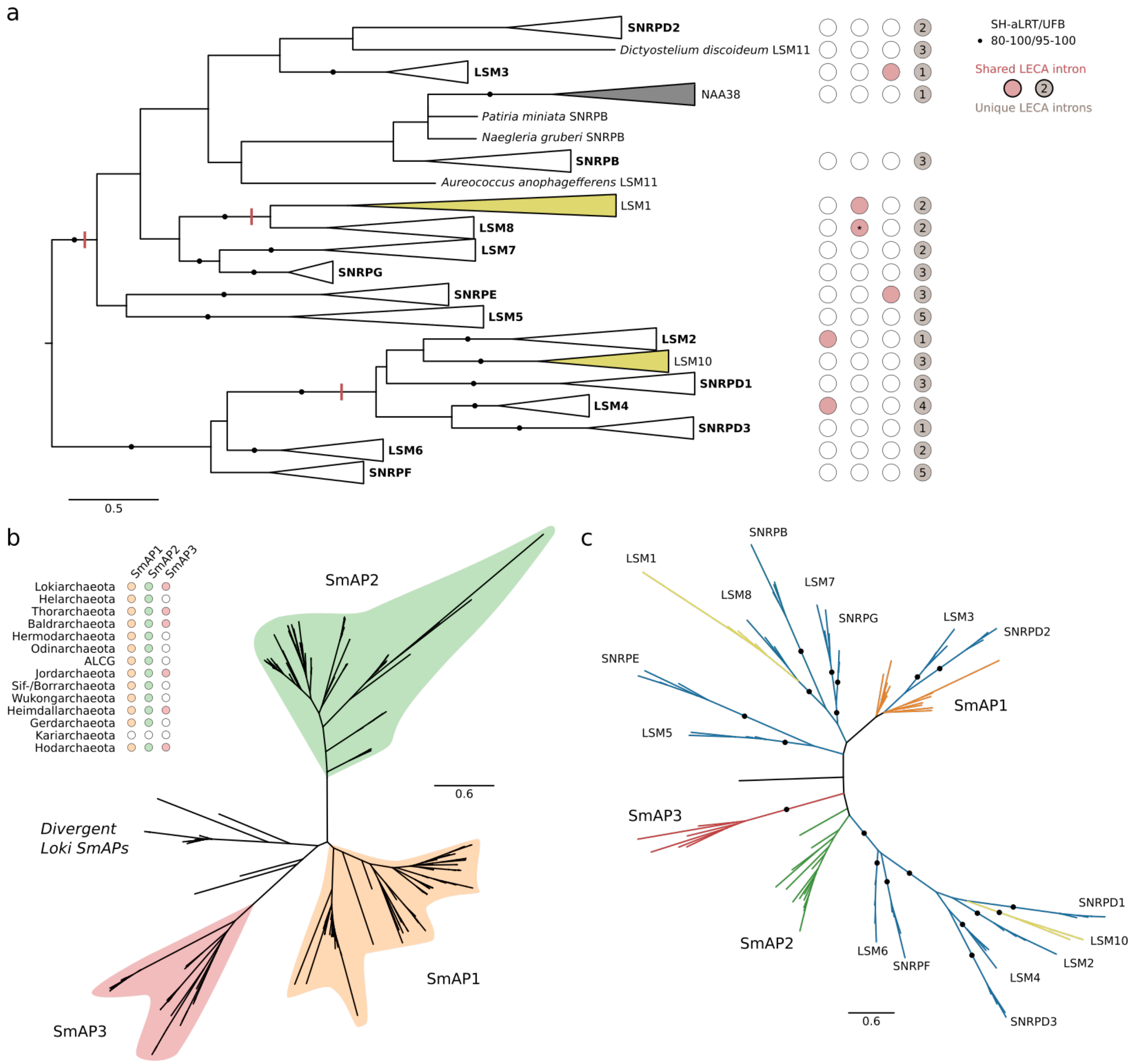


**Supplementary Figure 3. Evolutionary history of Lsm/Sm proteins.**
(a) Phylogeny of eukaryotic Lsm/Sm proteins. The tree was rooted using midpoint rooting. The intron positions that are shared between paralogs are indicated. The intron with an asterisk was classified as U12-type intron (Vosseberg et al. 2022). (b) Phylogeny of SmAPs in Asgard archaea. The names follow the classification used in previous work (Mura et al. 2003). The presence distribution shows that SmAP3 is restricted to a subset of taxa. (c) Phylogeny of eukaryotic and Asgard archaeal Sm-family proteins.

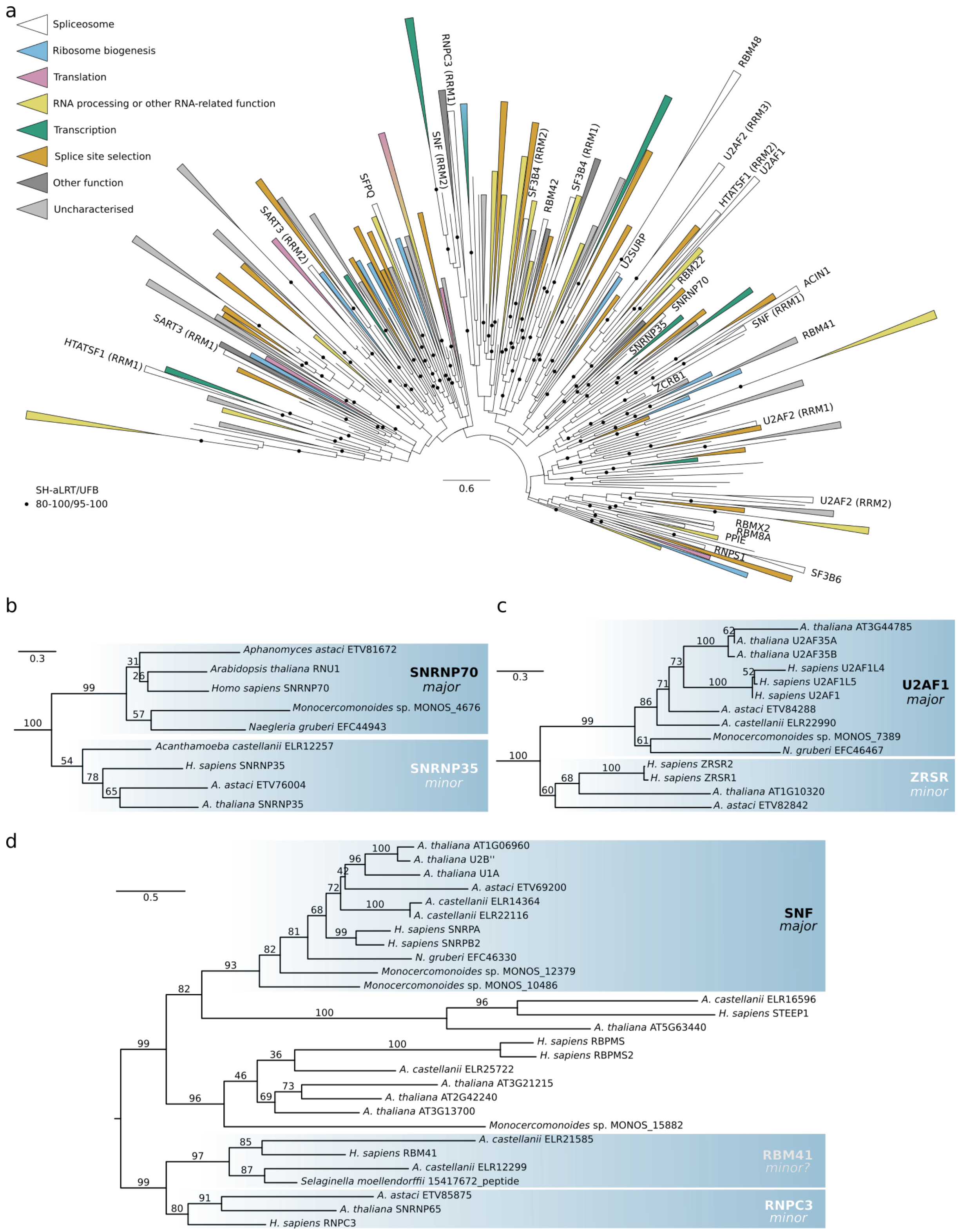


**Supplementary Figure 4. Evolutionary history of RRM proteins.**

(a) Phylogeny of RRMs. LECA OGs are collapsed and coloured based on their function. Names are only shown for the spliceosomal OGs. (b) Phylogeny of SNRNP70 and SNRNP35. The tree was rooted with PPIL4. (c) Phylogeny of U2AF1 and ZRSR. The tree was rooted with RBM39. (d) Phylogeny of SNF, STEEP1, RBPMS, RNPC3 and RBM41. The tree was rooted between SNF and RNPC3 based on the topology in the full tree. (b-d) Branch labels correspond with ultrafast bootstrap values. Major: major spliceosome-specific protein; minor: minor spliceosome-specific protein.

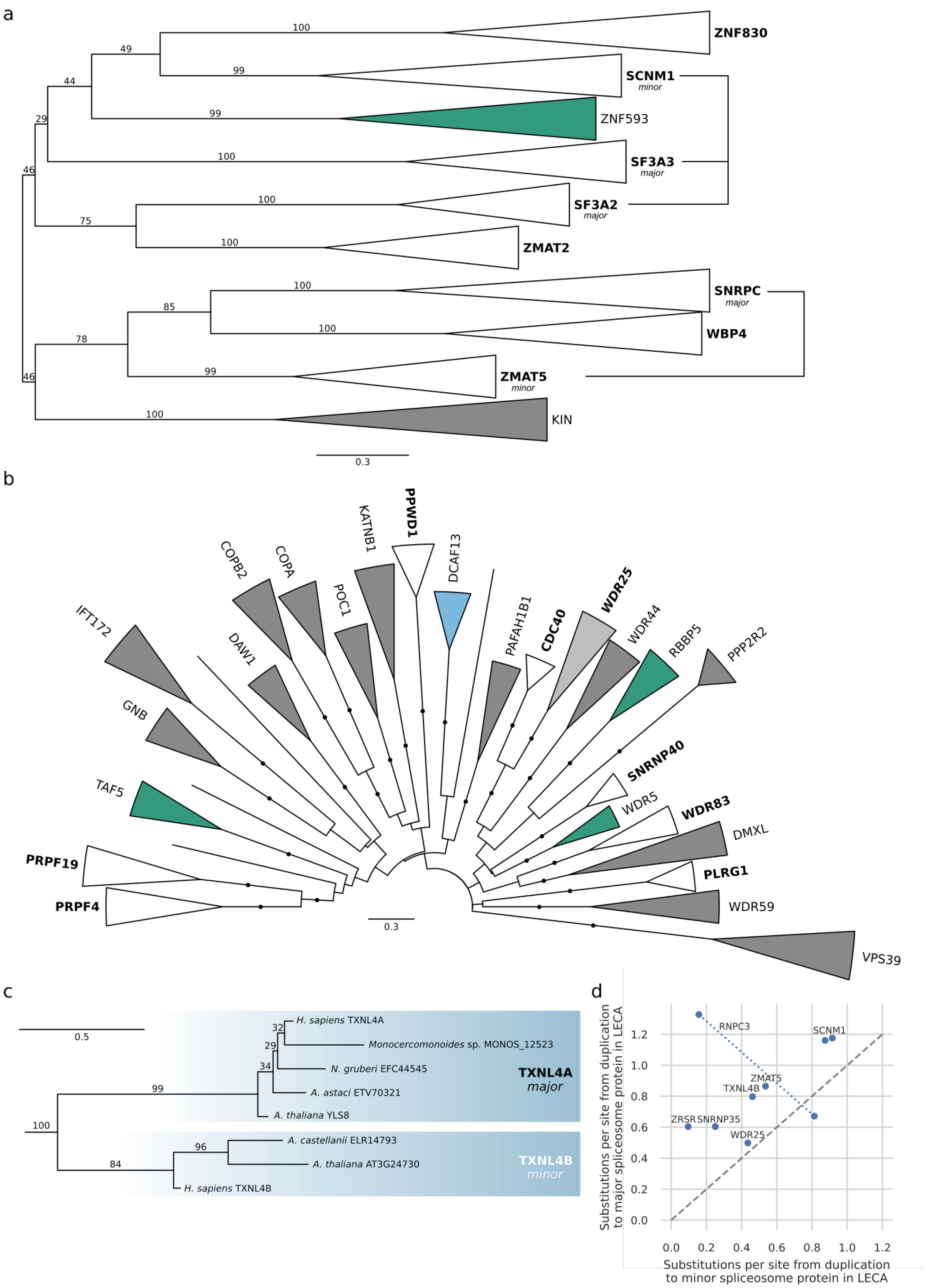


**Supplementary Figure 5. Evolutionary history of other large families that contributed to the spliceosome.** (a) Phylogeny of U1-type zinc finger proteins. (b) Phylogeny of spliceosomal and closely related WD40 proteins. (c) Phylogeny of TXNL4A and TXNL4B. The tree was rooted with NXN. (d) Relation between branch lengths of major and minor spliceosome-specific proteins. Because no outgroup was used for the SNF/RNPC3 root, the range of possible duplication-to-LECA branch lengths is plotted. (a, c) Branch labels correspond with ultrafast bootstrap values.

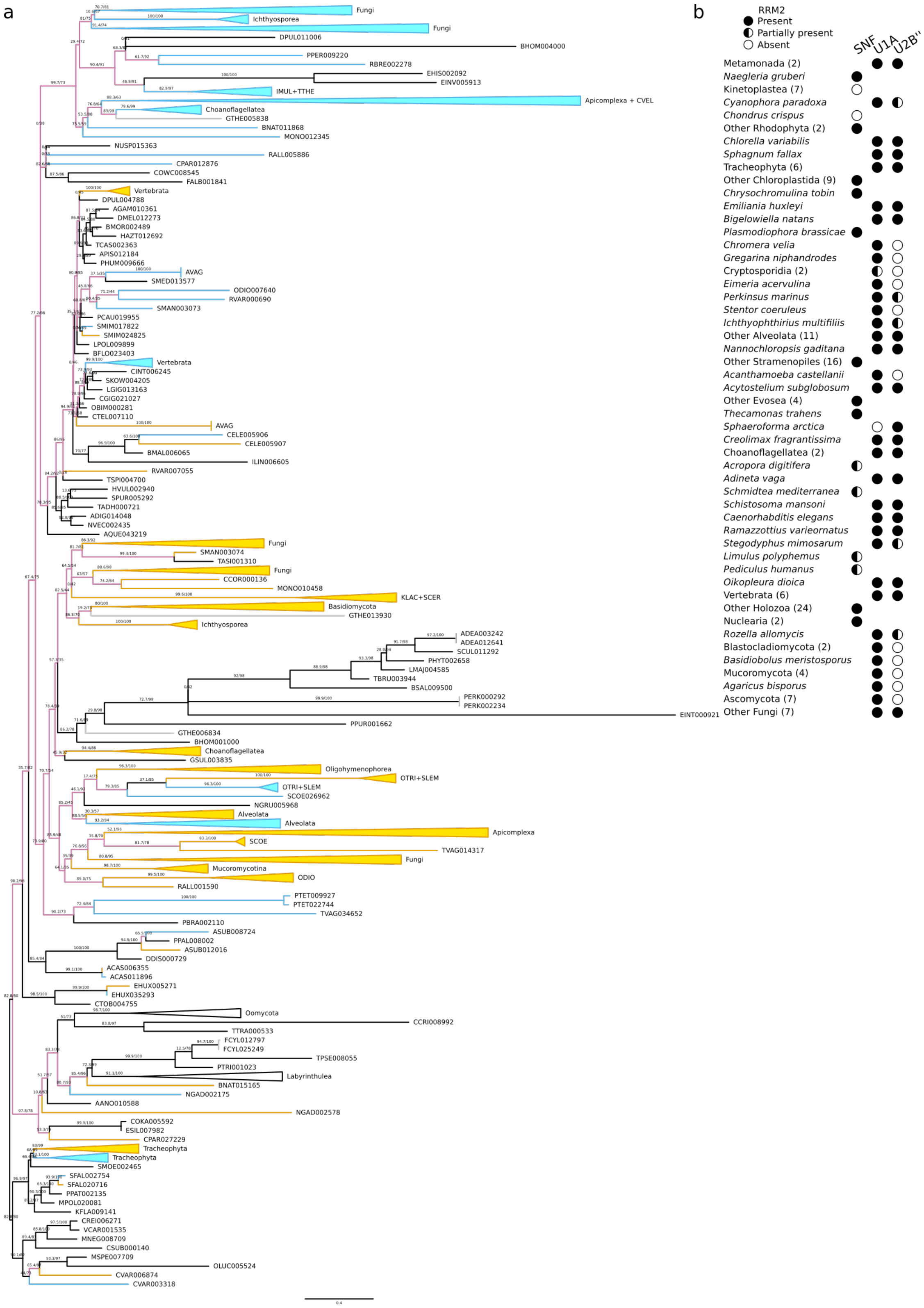


**Supplementary Figure 6. Evolutionary history of SNF proteins after LECA.**

(a) Phylogenetic tree of SNF. The four letters in the sequence identifiers refer to the species name (see Supplementary Table 1 in (Deutekom et al. 2019)). Paralogs are coloured based on their predicted fate, U1A (blue) or U2B'' (yellow), including adjustments. Pink branches connect U1A and U2B'' fate proteins of the same species, if necessary. Grey branches reflect duplications without a different fate prediction and the *G. theta* proteins. Branch labels correspond with the SH-like approximate likelihood ratios and ultrafast bootstrap values in percentages. (b) Presence profile of a second RRM domain in case of a single copy or two paralogs. Numbers in parentheses reflect the number of species.

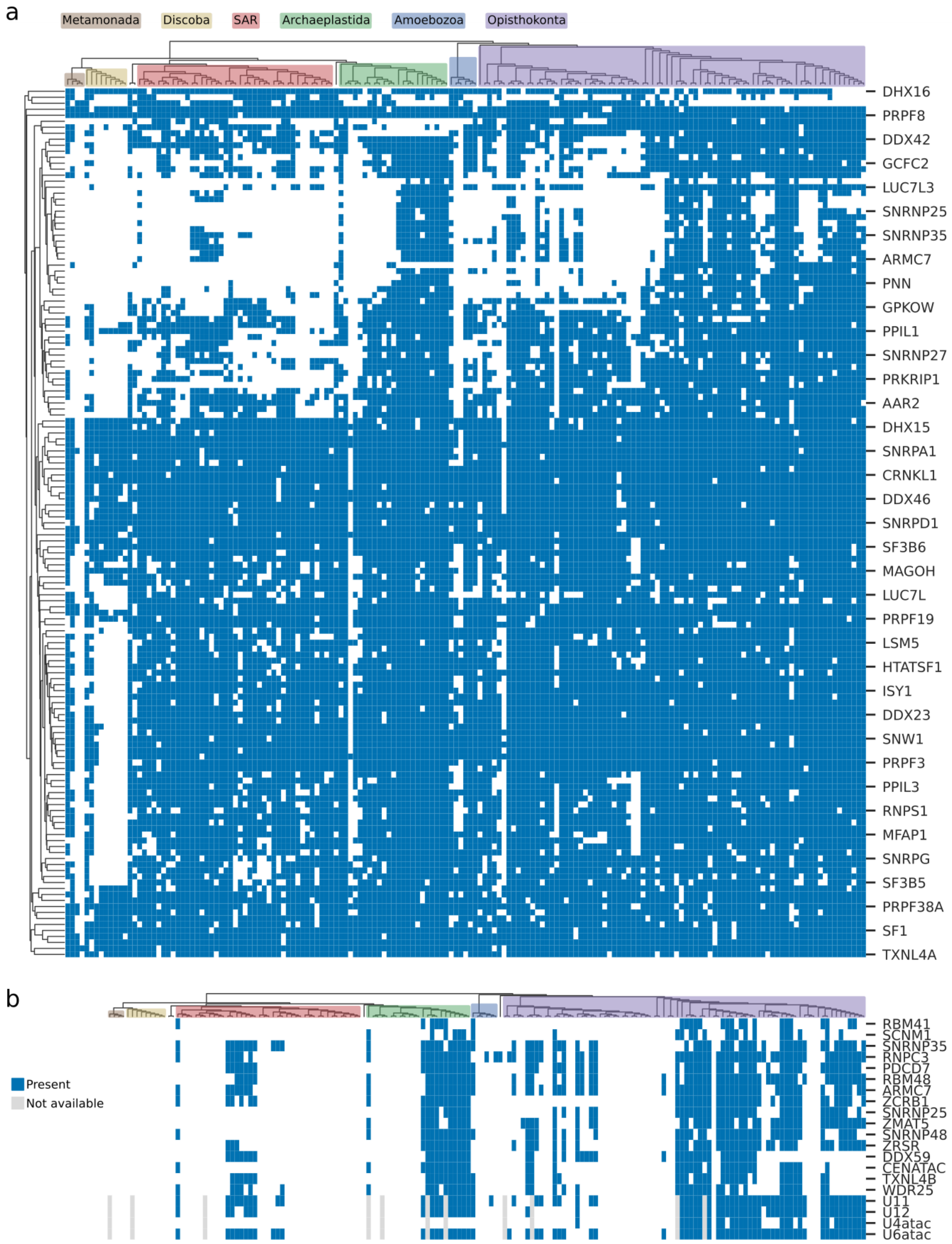


**Supplementary Figure 7. Presence of spliceosomal LECA OGs across eukaryotes.**
(a) Presence of the 145 spliceosomal LECA OGs across 167 eukaryotes. The columns correspond with the species, clustered based on the species tree. Eukaryotic groups are highlighted with different colours. The rows correspond with the spliceosomal LECA OGs; the names of some are indicated. The OGs are clustered with average linkage based on correlation distances. (b) Presence of minor-spliceosome specific proteins and snRNAs across eukaryotes. The candidate minor-spliceosome specific proteins RBM41, WDR25 and DDX59 are included.

**Supplementary Tables**

Supplementary Table 1. Spliceosomal proteins in human.

Supplementary Table 2. Spliceosomal proteins in baker's yeast.

Supplementary Table 3. Spliceosomal LECA OGs.

Supplementary Table 4. Evolutionary histories of spliceosomal LECA OGs.

Supplementary Table 5. Introns inferred in LECA for the spliceosomal LECA OGs and the numbers of intron positions shared with paralogs.

Supplementary Table 6. Programs and settings used for phylogenetic inference.
